# Rapid Invisible Frequency Tagging (RIFT) in a novel setup with EEG

**DOI:** 10.1101/2024.02.01.578462

**Authors:** Kabir Arora, Surya Gayet, J. Leon Kenemans, Stefan Van der Stigchel, Samson Chota

## Abstract

Steady-State Visual Evoked Potentials (SSVEPs) provide a report-free and continuous measure of neural processing. Recent progress in display technology has allowed for the tagging of multiple stimuli simultaneously at >60Hz frequencies - high enough to evade perceptibility, while still evoking an oscillatory neural response. Known as Rapid Invisible Frequency Tagging (RIFT), this technique has currently only been used in combination with Magnetoencephalography (MEG), which is less accessible compared to Electroencephalography (EEG). Although responses to LEDs flickering at similar frequencies have been shown in EEG, it is currently unclear whether RIFT, using a more conventional stimulus display, can sufficiently evoke a response in EEG, and therefore whether it is worth adding the RIFT-EEG pairing to the cognitive neuroscientist’s toolkit. Here, we successfully implement the first RIFT-EEG setup. We show that the oscillatory input is measurable in the EEG trace, what its topographical spread is, a rough range of applicable frequencies, and that this response is comparable to that evoked in MEG.

## 1 Introduction

Steady-State Visual Evoked Potentials (SSVEPs) are a characteristic neural response to rhythmically varying sensory inputs. As opposed to Event-Related Potentials (ERPs), which reveal responses to single, disconnected events, SSVEPs provide a continuous response to consistent visual or auditory stimuli directly from the neural traces without the need for manual report. This has led to their frequent use in understanding cognitive functions (See Norcia et al., 2015 for overview). An extensively used application of the SSVEP is frequency tagging, that is, labeling different stimuli with signature luminance oscillations that can later be disentangled and uniquely tracked over time from the neural response. Here, the SSVEP response to frequency tagged stimuli is used as a direct marker and tracker of spatial attention (Morgan et al., 1996; M. M. Müller & Hüb-ner, 2002) or feature-based attention (M. Müller et al., 2006; Pei et al., 2002). Recent progress in display technology has given rise to a new branch of frequency tagging: Rapid Invisible Frequency Tagging (RIFT), wherein stimuli are flickered at frequencies higher than 60Hz (Seijdel et al., 2023; Zhigalov et al., 2019).

This novel framework avoids two major limitations of existing SSVEP applications (which have mainly made use of stimuli flickering at rates below 30Hz). Firstly, at these frequencies, luminance changes are visible. This may confound effects of endogenous attentional shifts through exogenous flicker-caused distractions (Cass et al., 2011). RIFT, by flickering complex stimuli beyond the threshold of visibility, offers an attentional tracker without perceptual interference. Secondly, in the <30Hz range SSVEP responses may be difficult to disentangle from endogenous oscillations in similar frequency bands, or may even entrain or disrupt them (Notbohm et al., 2016; Spaak et al., 2014). In addition to operating far from the domain of alpha oscillations, RIFT has also been shown not to entrain endogenous oscillations in the gamma range (Duecker et al., 2021), circumventing any such confounds during analysis.

At this stage, such early studies have already answered certain fundamental questions about the utility of RIFT across various applications. It has been shown that RIFT is a powerful tool to capture spatial shifts of attention both to tagged stimuli (Bouwkamp et al., 2023, Preprint), as well as to tagged, but perceptually indistinguishable regions of visual space (Brickwedde et al., 2022). Over the last few years RIFT has been applied within the visual domains of reading (Pan et al., 2021), distractor suppression (Ferrante et al., 2023), and visual search (Spaak et al., 2023, Preprint) among others, establishing that its robustness is comparable to that of the more conventional SSVEP. Current RIFT work is expanding both upon the cognitive contexts that may benefit from the technique (Bouwkamp et al., 2023; Seijdel et al., 2024, Preprint), as well as technical aspects such as optimizing display features (Minarik et al., 2023) and exploring alternate forms of tagging (Spaak et al., 2023, Preprint).

Despite the clear and promising potential such work shows for RIFT within vision research, relatively few studies have utilized it. One cause may be the accessibility of current RIFT setups. Beyond the display equipment itself, all existing RIFT research has been implemented in combination with MEG. While the existing framework has clearly delivered novel insights for cognitive neuroscience as described above, the low number of MEG setups worldwide combined with both the active cost of running MEG experiments as well as the initial capital required encourages the exploration of viable, more accessible alternatives. Additionally, as the potential of RIFT as a communication medium for Brain-Computer Interfaces begins to be explored (Brickwedde et al., 2022) as an improvement upon existing SSVEP Brain-Computer Interfaces (Zhu et al., 2010), the need to assess the possibility of a more portable RIFT framework rises further.

Electroencephalography (EEG) has been shown to be sensitive to periodic stimulation in the RIFT frequency range using flickering LEDs (Gulbinaite et al., 2019; Herrmann, 2001). However, most tasks designed to study cognition utilize stimuli varying across several features such as size, shape, orientation, colour, etc. and therefore flickering LEDs may not reflect the context within which RIFT could be best utilized. Here, we successfully implement the first RIFT setup paired with EEG. We show that it is possible to pick up RIFT signals from multiple stimuli at different frequencies in the EEG response, reveal the topographical spread of these signals, and demonstrate that the evoked signals are comparable in magnitude to those evoked in MEG.

## 2 Methods

### Participants

We recruited 36 healthy participants (29 female, 23.3 *±* 3.0 | mean *±* std) with normal or corrected to normal vision. None of the participants reported a history of epilepsy or psychiatric diagnosis. Participants were compensated either with €20 or an equivalent amount of participation credits as per Utrecht University’s internal participation framework. The study was carried out in accordance with the protocol approved by the Faculty of Social and Behavioural Sciences Ethics Committee of Utrecht University.

### Experimental Design

Trials began with a centrally presented fixation cross (uniformly random duration between 1-1.25s), after which both memory stimuli were presented (1s). This was followed by a variable stimuli-cue delay (uniformly random, 1.5-1.9s), at the end of which a central retrocue was presented (0.15s) in the form of either a blue or red circle, informing participants which item would be probed on that trial. The retro-cue randomly cued either the left or the right item (50/50). After a fixed cue-probe interval (1.5s), the memory probe was displayed centrally until response. Participants had to specify with a keyboard button press whether the orientation of the memory probe was more clockwise (Q key) or anti-clockwise (P key) compared to that of the retro-cued memory stimulus. Participants were able to respond 0.5s after the memory probe onset (indicated to them via a change in the colour of the fixation cross from black to green). If no response was received within 3s, or a different button was pressed, the trial ended and participants were shown ‘incorrect’ feedback (median = 0.4% of trials).

Each participant completed 480 trials (excluding 10 practice, not analyzed; 15 blocks of 32 trials each). Orientations of the memory stimulus (always distinct), as well as whether the blue/red stimuli would be presented to the left or right, were equated in prevalence individually and presented in random order.

### Stimuli

The screen background was maintained at true grey throughout the experiment. A black fixation cross (0.4 dva) was present in the center of the screen throughout each trial. Memory stimuli consisted of circular square wave gratings (*r* = 3 dva, spatial freq. = 2 cpdva) either blue or red in colour. These were presented slightly below horizontal (eccentricity = 6 dva horizontal, -2 dva vertical) in order to facilitate the tagging response (Minarik et al., 2023). To minimize tagging frequency, a ra--dially symmetric transparency mask was applied to the memory stimuli as per the following function:

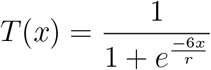

where *x* is the distance of a point on the circular patch from center (0) to the circumference (at radius *r*), and *T* (*x*) is the resulting transparency at this point ranging from 0.5 (semi-transparent) to 1 (fully opaque). Both memory stimuli were presented with distinct orientations ranging from 0-165° (in intervals of 15°). The retro-cue (*r* = 0.23 dva) was displayed centrally in one of the two memory stimuli colours. The memory probe consisted of a black, circular ring (*r* = 1.2 dva, thickness = 0.13 dva) with spokes (0.6 dva extensions) protruding outwards from diametrically opposite ends to indicate an orientation. These dimensions ensured that the tips of the memory probe never overlap (minimum separation = 1.5 dva) with the flickering regions on screen (See *Tagging Manipulation*). The memory probe angle varied between 3°-50° clockwise or anti-clockwise of the retro-cued memory stimulus, following a staircase (Farell & Pelli, 1999) that maintained accuracy on the memory task at 75% (PsychtoolBox QUEST algorithm, *β* = 3.5, Δ = 0.01, *γ* = 0.5). Feedback was given after each trial in the form of a green/red image (*r* = 1.5 dva) of a tick/cross for right/wrong answers respectively.

### Tagging Manipulation

The areas corresponding to the two memory stimuli were tagged from stimuli onset until the end of the trial. Two frequencies were randomly assigned to either the left or right area. The two frequencies used were either 60/64Hz (24 participants) or 60/68.57Hz (12 participants). The lateralization of the frequencies (60L/60+R vs. 60+/60R) was counterbalanced with the direction of the retro-cued item and presented in a random sequence. For the first 1s of a trial (when memory stimuli were being presented), the two circular gratings were tagged with the corresponding frequency. For the remainder of the trial, the background was tagged (range 0-1 luminance in order to look invisible against the true grey background). The same transparency map was applied to the flickering regions as the memory stimuli (See *Stimuli*). The tagging sinusoids were constructed such that despite the variable stimuli-cue delay, the waves were always at the same phase at the moment of retro-cue onset. Temporal precision of the displayed stimuli was continually recorded using PsychToolBox’s Screen(‘Flip’) command. Any trial with a frame displayed >4ms off-time was excluded from analysis (mean = 0.17% of trials, median = 0 trials).

### Protocol

Participants underwent a 2-hour session at the Division of Experimental Psychology, Utrecht University. Participants received procedural information prior to the session, and provided informed consent, date of birth, biological sex, and dominant hand information at the beginning of the session. After completion of the EEG setup, participants were seated 76cm from the screen with a chinrest. After eye calibration, the experiment was explained using a visual guide and verbal script. Participants were also informed that they may potentially see visual glitches or flickers on the screen (due to the high-speed projection), and that they would be asked after the experiment whether they did or not. Following these instructions and 10 practice trials, the 1 hour experiment was completed. Participants filled out a questionnaire on whether they noticed any visual artifacts on screen (and if so, at what stage of the task, and to what degree they felt this interfered with their task on a scale of 1-5). Compensation was awarded when applicable, and the session was ended.

### Display Apparatus

Stimuli were projected using a ProPixx projector (VPixx Technologies Inc., QC Canada; resolution = 960×540px; refresh rate = 480Hz) in a rear-projection format (projected screen size = 48×27.2cm). Experimental code was written using PsychToolBox (Brainard & Vision, 1997; Kleiner et al., 2007) in MATLAB (MATLAB, 2022).

### Eye-tracking Recording and Analysis

Gaze was tracked using an Eyelink SR (SR Research, Ontario, Canada) eyetracker. Both eyes were tracked at 500Hz. Immediately prior to the experiment, a 9-point calibration was performed. This calibration was also performed after every 3rd block. Eye position data was not analyzed in the current study.

### EEG Recording and Pre-processing

EEG data was recorded using a 64-channel ActiveTwo BioSemi system (BioSemi B.V., Amsterdam, The Netherlands) at 2048Hz. To record vertical and horizontal eye movements, two additional electrodes were placed above and on the outer canthus of the left eye respectively. Immediately prior to the experiment, adequate signal quality from all channels was ensured using BioSemi ActiView software. All data analysis was conducted in Matlab using the Fieldtrip toolbox (Oostenveld et al., 2011). The EEG data was first re-referenced to the average of all channels (excluding poor channels determined by visual inspection, median = 5 [frontal] channels per participant). Data was high-pass filtered (0.01Hz), then line noise and harmonics were removed using a DFT filter (50, 100, 150Hz). Data was segmented into trials ranging from 3.4s before to 2s after retrocue onset. An ICA was performed to remove oculomotor artifacts, and trials with other motor artifacts (as per visual inspection) were removed. Baseline correction was performed using a 0.5s window (−0.7 to -0.2s w.r.t stimuli onset).

### RIFT Response: Coherence

In order to determine the strength of the EEG response to RIFT frequencies, magnitude-squared coherence was used, which is a dimensionless quantity from 0-1 that measures how consistently two signals are matched in both their frequency content and phase. This results in higher values when two signals oscillate at the same frequency while doing so with a constant phase-difference across trials. Coherence was computed between a reference wave (pure sinusoids at the corresponding 60Hz/64Hz/68.57Hz frequency, sampled at 2048Hz) and sets of trials per channel and participant. The 5.3s trials were first band-pass filtered (*±*1.9Hz) at the frequencies of interest (60Hz & 64Hz for the first set of participants, 60Hz & 68.57Hz for the second set) using a two-pass Butterworth filter (4th order, hamming taper). The filtered time-series data was Hilbert transformed. This provided a time-varying instantaneous magnitude (*M*(*t*)) and phase (*ϕ*(*t*)). The set of all instantaneous magnitudes of the filtered responses (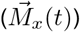) and the reference sinusoid (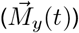) across all *n* trials, as well as the differences between their instantaneous phases across all *n* trials (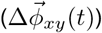) were used to compute time-varying coherence:

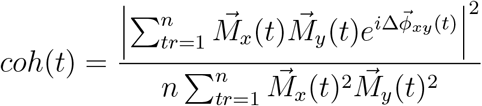

In order to compute coherence spectrograms and inspect the sharpness of the coherence signal, coherence was computed in a similar manner for frequencies ranging from 56.5Hz to 71.5Hz in intervals of 0.5 Hz.

### RIFT Response: Power

For visual comparison, the strength of the RIFT response was also computed through Fourier Transforms. An FFT was applied to each 5.3s trial using a hamming window. Signal-to-noise (SNR) was then computed by dividing power at the frequencies of interest (60Hz, 64Hz, and 68.57Hz) with the mean power of surrounding frequencies excluding close neighbours (*±*(0.5-2Hz) away).

## 3 Results

We first assessed whether the RIFT signal emerges within the EEG response at all. Time-varying coherence measures were averaged across all experimental conditions and participants to provide a general overview. For this initial outline, six channels with the highest mean coherence were identified per participant and averaged. Data from the first set of participants (using 60 and 64Hz, n=24) and the second (using 60 and 68.57Hz, n=12) are presented separately. The computed coherence spectrograms show that for both groups of participants (Figure 3.1a-b) there are notable peaks in oscillatory activity at the tagged frequencies. A spread within the frequency domain is also present and equivalent to that of the bandpass filter range (*±* 1.9Hz) applied prior to applying the coherence measure. We then inspected the spatial spread of this obtained response over the EEG topography averaged across the entire flicker duration (Figure 3.2a-b) both with coherence and power/SNR. Both topographies are restricted to parietal/occipital electrodes.

**Fig 2.1:**
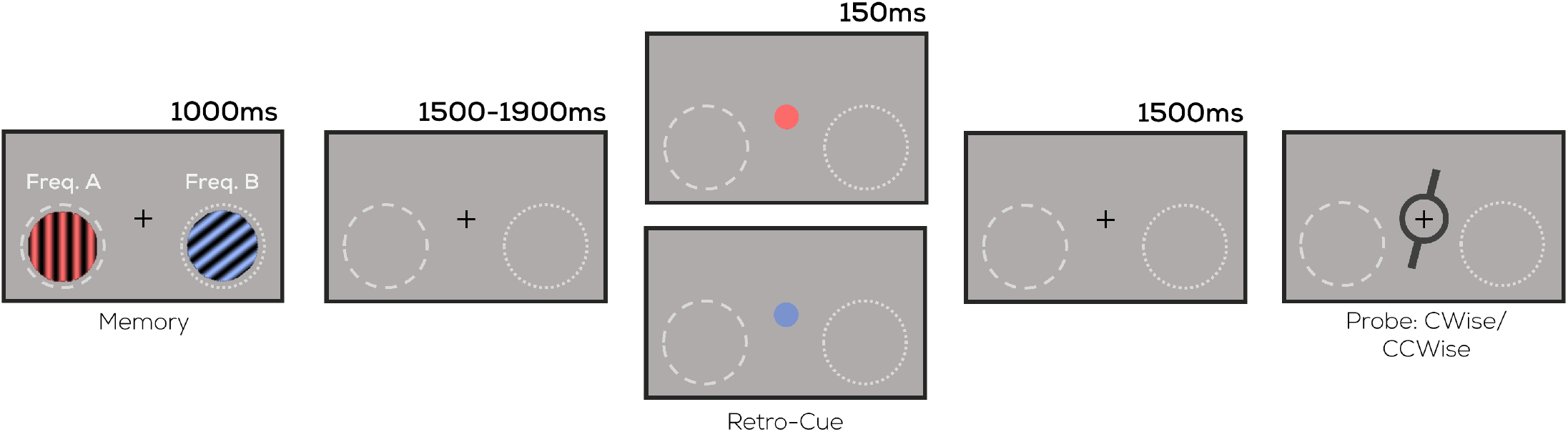
Task design. Two uniquely coloured and oriented stimuli were presented on either side of fixation. A colour-based retro-cue specified which of the two would be probed for response. Participants indicated whether the orientation of a memory probe was more clockwise or counterclockwise compared to that of the retro-cued item. The highlighted locations were flickered at 60/64/68.57Hz (See Tagging Manipulation) throughout the trial (Note: not to scale; memory probe did not overlap with flickering regions in actual display).

**Fig 3.1:**
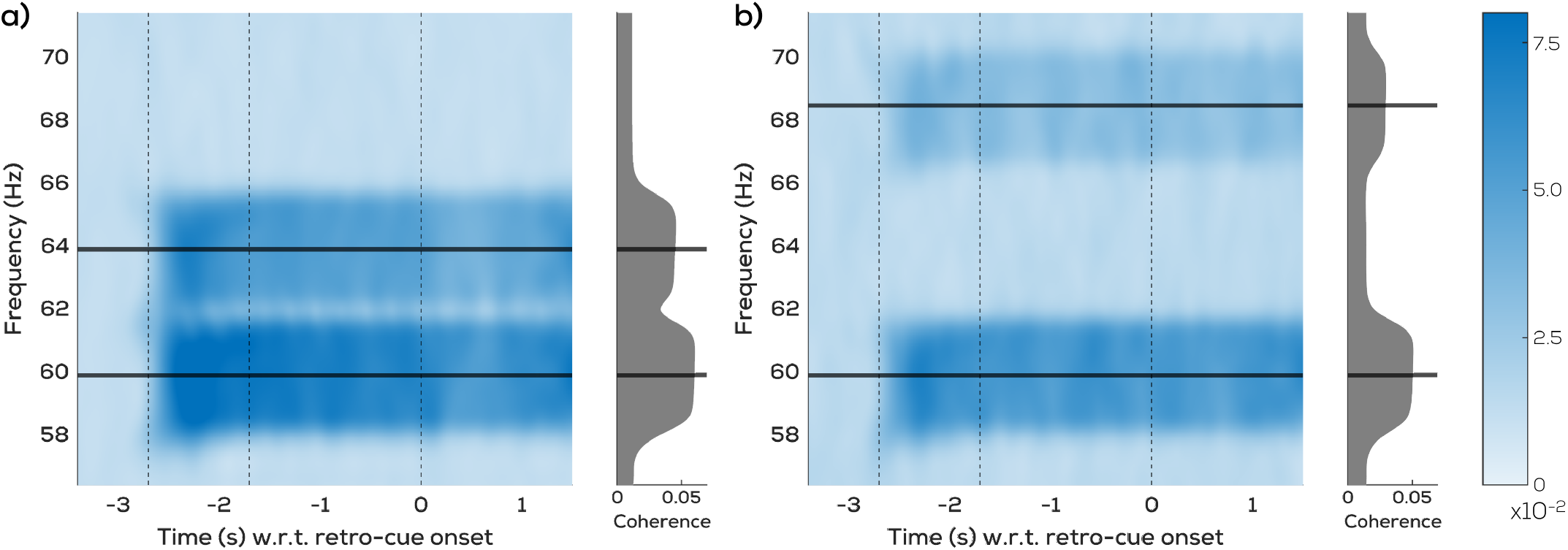
Coherence Spectrogram. Mean coherence (averaged across participants and best channels, see text) for **a)** Group 1 participants with tagging at 60 & 64Hz (n=24) **b)** Group 2 participants with tagging at 60 & 68.57Hz (n=12). Dotted lines indicate stimuli onset, stimuli offset, and retro-cue onset respectively.

**Fig 3.2:**
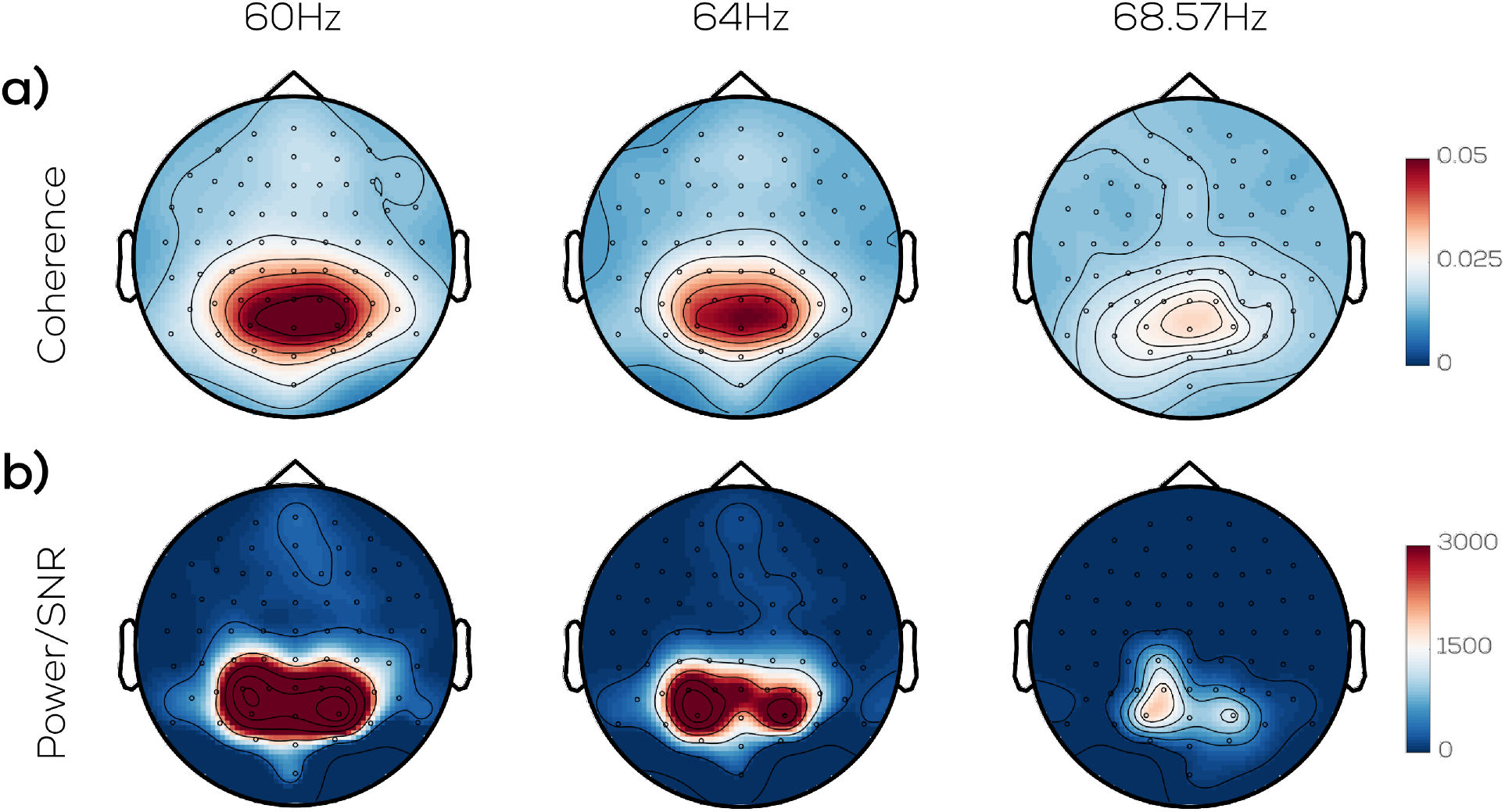
RIFT topography in EEG, using **a)** Coherence **b)** Power/SNR at 60Hz (n=36), 64Hz (n=24), and 68.57Hz (n=12), averaged across participants and experimental conditions.

Given the clear drop in signal evoked from 60Hz and 64Hz to 68.57Hz (Figure 3.2),we next determined quantitatively whether these tagging frequencies implemented in the present study were all equivalent options for RIFT-EEG in terms of the coherence elicited (Figure 3.3).

For this, per-participant we averaged the 6 channels with the strongest RIFT response at each frequency. Although no significant difference was seen across 60Hz and 64Hz (Mean difference = 0.0187; 95% confidence intervals of bootstrapped mean differences: [-0.0083, 0.0441]; *p*=0.19), the coherence elicited by 68.57Hz was significantly lower than that of 64Hz (Mean difference = -0.031; 95% CI: [-0.055, -0.008]; *p*<0.05) and that of 60Hz (Mean difference = -0.049; 95% CI: [-0.070, -0.028]; *p*<0.005). This is also reflected at the topography level. Visual inspection of coherence topographies for all participants at their respective tagging frequencies (Figure 3.4) showcases a clear parietal/occipital focus in most topographies at 60Hz and 64Hz. This is absent when inspecting the neural response at 68.57Hz.

**Fig 3:3.**
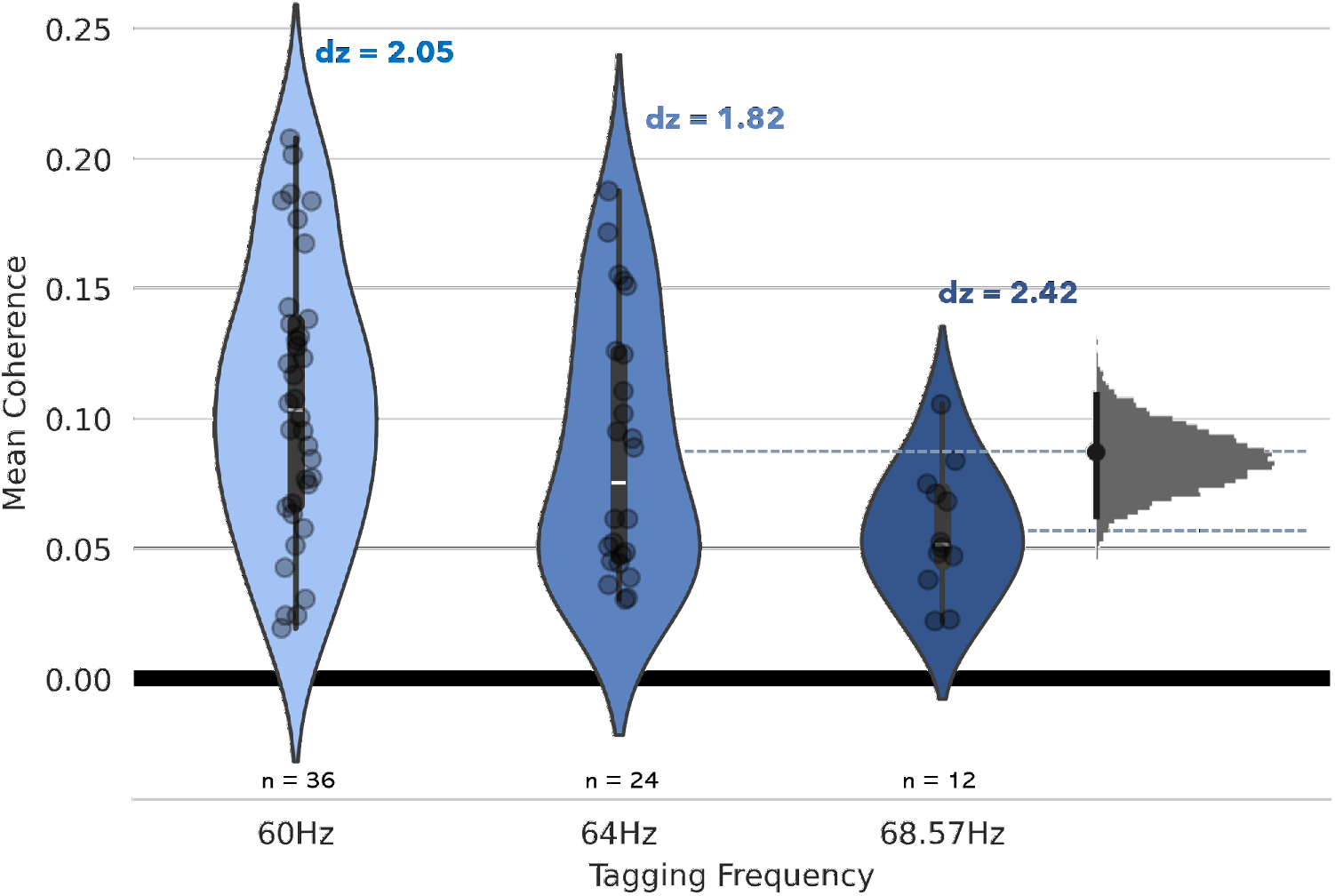
Mean coherence across tagged frequencies. (from top-6-channel averages per participant), averaged over tagged duration. Grey indicates distribution of bootstrapped mean differences between 64Hz and 68.57Hz (dashed lines indicate respective means; black bar indicates 95% CIs; other comparisons stated in text).

**Fig 3.4.**
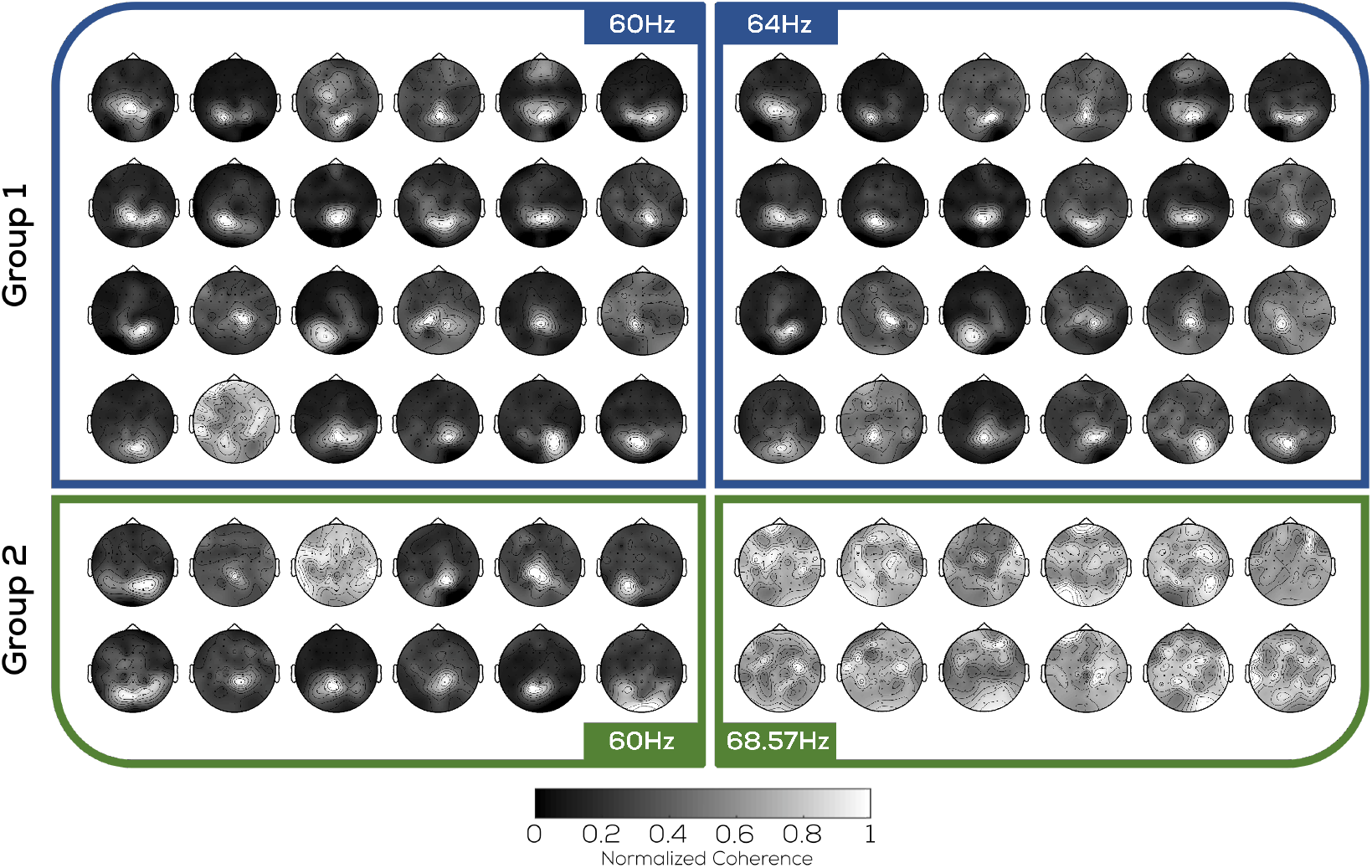
RIFT topography variability. Mean coherence across tagged frequencies and participants (normalized w.r.t. each participant’s most strongly responsive channel), averaged across tagged duration.

Lastly, we assessed the extent to which the RIFT response obtained through EEG is comparable to that of MEG (Figure 3.5). We compared the mean coherence elicited for all 36 participants in this study at 60Hz (from the 1s interval where stimuli were flickering on-screen) to equivalent values from a RIFT-MEG study on reading (Pan et al., 2021) after roughly adjusting for differences in flickering area by computing an evoked coherence value per dva2 of flickering area on screen. No significant difference was seen between MEG and EEG coherence per unit area (Mean difference = 0.0008; 95% CIs: [-0.0002, 0.0022]; *p*=0.19). The latter showed lower variance (EEG *µ*/*σ* =1.95, MEG *µ*/*σ* = 1.62).

Participants were also informed at the start of the experiment that they would be asked at the end whether they observed any visual artifacts on screen, and filled out a questionnaire on the subject at the end of the experiment. Of the 36 participants, only 4 responded positively to noticing any ‘flickering glitches’ or similar on the screen. Three of these reported only noticing such artifacts when the stimuli were on screen (and not during the stimuli-cue intervals), all reporting flicker intensity with the lowest possible score of 1/5. One participant reported only noticing such artifacts during the stimuli-cue intervals, with an intensity score of 2/5.

**Fig 3.5:**
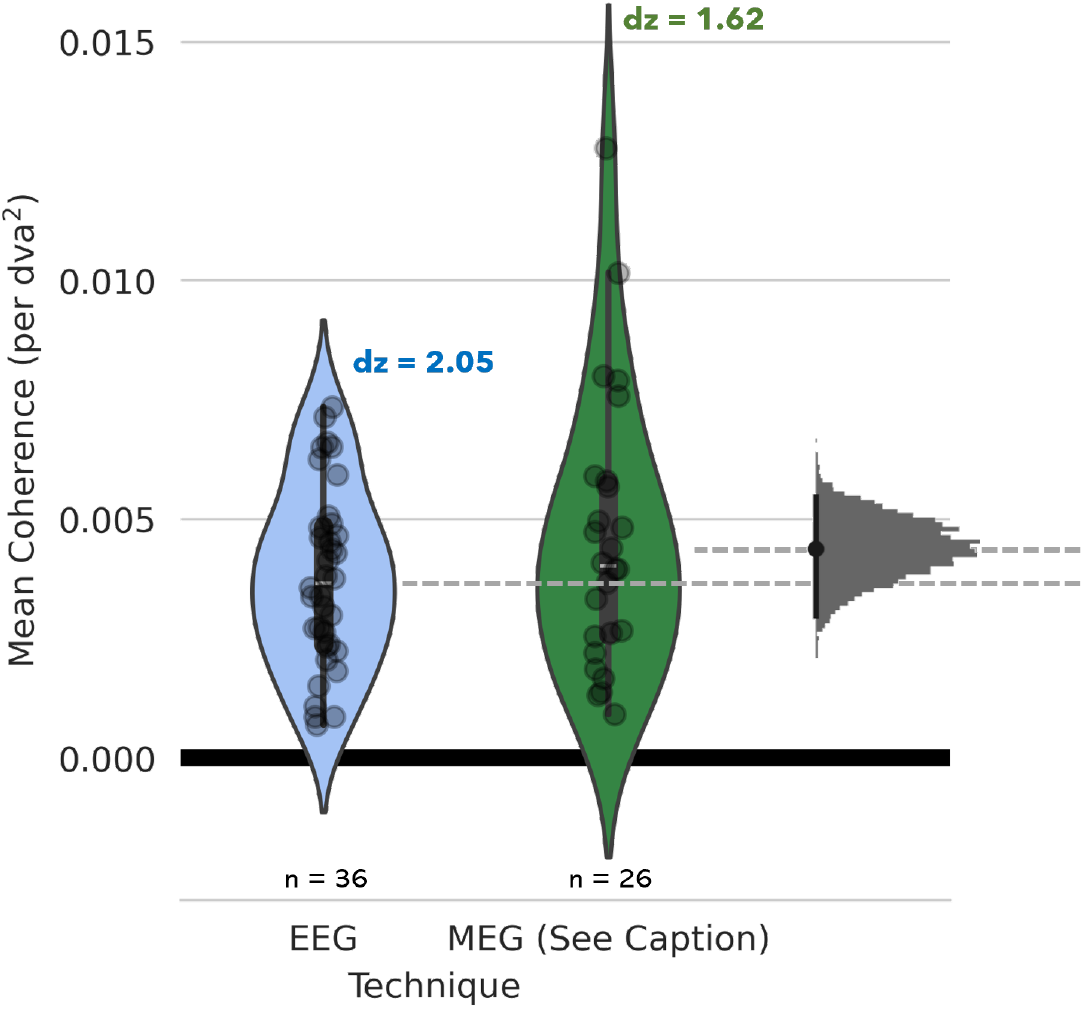
Coherence from different tools. Mean coherence at 60Hz across imaging modalities EEG (present study; from top-6-channel averages per participant) and MEG (Pan et al., 2021) averaged over tagged duration. Grey indicates distribution of bootstrapped mean differences between 64Hz and 68.57Hz; black bar indicates 95% CIs.

## 4 Discussion

We conducted a visual memory task with 36 participants in order to determine the feasibility of pairing RIFT with EEG. We show the frequency content of the EEG response to tagged stimuli/regions, the topographical spread of this response, cross-participant variations, and that it is comparable to that evoked in MEG.

The utility of RIFT in combination with EEG for cognitive neuroscience experiments may be qualitatively assessed (and compared to its utility with MEG) across two metrics. Firstly, the strength of the RIFT signal evoked in the neural response determines whether EEG is sensitive enough to pick up on such high-frequency oscillatory responses from conventional stimulus displays at all. Secondly, the range of frequencies available for tagging is relevant, since this directly impacts the number of stimuli that are ‘taggable’ and thus the possible types of experiments that may be compatible with an EEG-based RIFT setup.

### Overall RIFT-Response

Time-frequency analysis showed that there is an increase in oscillatory activity selectively at tagged frequencies upon presentation of luminance-modulated stimuli, and additionally that this increase is sustained even throughout periods where regions of space are tagged in the absence of a stimulus. We also confirm that this ’absence of a stimulus’ in RIFT-tagged regions is truly reflected, since almost none of our participants report noticing any flickers even though they were informed beforehand to expect such visual artifacts. We show that the signal emerges from parietal/occipital electrodes, visible both with more conventional FFT-based analysis as well as by using coherence measures across trials (with the latter providing a broader topography). The evoked strength of the RIFT-EEG response at 60Hz is similar to that in MEG, and there is proportionally similar cross-participant variance between the two, indicating comparable consistency. It is therefore clear that RIFT does in fact evoke a recoverable oscillatory response in EEG.

Importantly, the results presented here may actually not reflect the upper limit of RIFT strength that can be extracted from the EEG response; certain factors may, through alternative decisions of experimental design and analysis, evoke an even stronger response. Firstly, recent work with MEG has shown that accounting for minute equipment-driven phase-variability in the tagging signal across trials can improve signal strength (Spaak et al., 2023, Preprint). Secondly, the task we present here flickered stimuli at a higher eccentricity than the one we draw a comparison to (Pan et al., 2021), and it has been shown that RIFT signal strength varies inversely with flicker eccentricity in MEG (Minarik et al., 2023). This suggests that future optimization of methods may give rise to a stronger reflection of the tagged input in the EEG response.

### Dependence on tagged frequency

This study utilized three frequencies, namely 60Hz, 64Hz, and 68.57Hz, due to their compatibility with the ProPixx projector refresh rate. Although responses to flickering LEDs have been shown in EEG up until 80Hz (Gulbinaite et al., 2019) or more (Herrmann, 2001), our results show that there is a significant drop-off in RIFT signal acquisition at 68.57Hz. From the participant-wise topographies, it can be seen that plotting the RIFT coherence at 68.57Hz over the head topology does not evoke a stronger signal from parietal/occipital channels compared to the rest, in stark contrast to both 60Hz coherence topologies in the same participants and 60/64Hz topologies in another set of participants. Existing RIFT studies with MEG have utilized similar frequencies to equivalently tag three objects simultaneously (Bouwkamp et al., 2023, Preprint). Therefore, the range of frequencies that display an equivalent neural tagging response do not boast a similar robustness in EEG to that of MEG.

Though it may be possible to evoke a stronger RIFT response from higher frequencies in future tasks as described above, currently we only see usable responses to 60Hz and 64Hz tagging. However, given that previous RIFT work has also made use of 56Hz (Brickwedde et al., 2022) without any resulting concerns over perceptibility of the flicker, and that different choices during the analysis stage (narrower bandpass filtering for coherence) may allow increased frequency resolution for tagging, we infer that RIFT-EEG is capable of uniquely tagging at least two individual stimuli/regions, with a strong likelihood of being able to tag at least three.

We conclude that these metrics offer sufficient evidence in support of RIFT-EEG as a tool sensitive enough to reflect modulations from shifts of attention. This, in combination with developments towards accessibility on the display side of RIFT, such as commercially available monitors with 500Hz refresh rates (Alienware Gaming Monitor, Dell Inc., Texas) exceeding the 480Hz refresh used here, suggests a promising future for the widespread application of RIFT to understanding cognitive functions as well as to Brain-Computer Interfaces.

## Acknowledgements

This project receives funding from the European Research Council (ERC) under the European Union’s Horizon 2020 research and innovation programme (grant agreement n° 863732). The authors thank Katharina Duecker for valuable feedback on our experimental setup and analysis pipeline, and Laura van Zantwijk for assistance with data collection.

## Notes

### Competing Interest Statement

The authors have declared no competing interest.

